# A Photonic Resonator Interferometric Scattering Microscope for Label-free Detection of Nanometer-Scale Objects with Digital Precision in Point-of-Use Environments

**DOI:** 10.1101/2022.12.13.520266

**Authors:** Leyang Liu, Joseph Tibbs, Nantao Li, Amanda Bacon, Skye Shepherd, Hankeun Lee, Neha Chauhan, Utkan Demirci, Xing Wang, Brian T. Cunningham

## Abstract

Label-free detection and digital counting of nanometer-scaled objects such as nanoparticles, viruses, extracellular vesicles, and protein molecules enable a wide range of applications in cancer diagnostics, pathogen detection, and life science research. The contrast of interferometric scattering microscopy is amplified through a photonic crystal surface, upon which scattered light from an object combines with illumination from a monochromatic plane wave source. The use of a photonic crystal substrate for interference scattering microscopy results in reduced requirements for high-intensity lasers or oil-immersion objectives, thus opening a pathway toward instruments that are more suitable for environments outside the optics laboratory. Here, we report the design, implementation, and characterization of a compact Photonic Resonator Interferometric Scattering Microscope (PRISM) designed for point-of-use environments and applications. The instrument incorporates two innovative elements that facilitate operation on a desktop in ordinary laboratory environments by users that do not have optics expertise. First, because scattering microscopes are extremely sensitive to vibration, we incorporated an inexpensive but effective solution of suspending the instrument’s main components from a rigid metal framework using elastic bands, resulting in an average of 28.7 dBV reduction in vibration amplitude compared to an office desk. Second, an automated focusing module based on the principle of total internal reflection maintains the stability of image contrast over time and spatial position, facilitating automated data collection. In this work, we characterize the system’s performance by measuring the contrast from gold nanoparticles with diameters in the 10-40 nm range and by observing various biological analytes, including HIV virus, SARS-CoV-2 virus, exosomes, and ferritin protein.

## 1. Introduction

The ability to detect and characterize biological nano-objects at digital resolution without extrinsic labels or labor-intensive assay preparation is the core advantage of interferometric scattering microscopy (iSCAT). Since its introduction nearly two decades ago [1], iSCAT has attracted significant interest in the field of bioimaging applications, including biomolecular and nanomaterial mass photometry [2-4], protein interaction and oligomerization characterization [5, 6], three-dimensional particle tracking, and nanoparticle classification [7, 8]. Imaging contrast in iSCAT relies on the fact that the detected signal is linearly dependent upon the volume of the scattering particle rather than the volume squared, as for pure scattering-based microscopies. The linearity is a result of the interference cross-term between the object-scattered light and the background reference light [9]. Consequently, by carefully manipulating the intensity ratio between the two light sources, one could push the detection limit for protein molecules to molecular weights as small as 65 kDa [10]. One way to enhance the signal contrast is by attenuating the overwhelming reference light intensity through methods such as thin film interference [11], back focal plane engineering [12], and polarization filtering [13]. Alternatively, the scattered light can be strengthened to increase the magnitude of the detected signal. For example, surface plasmon resonance has been employed in conjunction with iSCAT to improve the ratio between the scattered light intensity and the background [14]. Similarly, metallic nanoparticles exhibit an enhanced scattering cross section under localized surface plasmon resonance conditions [15], enabling their use as labels for biosensing applications with iSCAT instruments.

Recently, we demonstrated the concept of utilizing a resonantly reflective photonic crystal (PC) metamaterial surface in combination with iSCAT to enhance scattered field intensity while also reducing the magnitude of background reference light intensity that interferes with scattered light from the detected nano-object [16], in an approach called Photonic Resonator Interferometric Scattering Microscopy (PRISM). In PRISM, the PC periodic structure is resonant with the wavelength of the laser illumination, resulting in the formation of electromagnetic standing waves in the form of surface-confined evanescent fields. As the electric field magnitude of the standing waves is magnified with respect to the electric field of the illumination source, we showed that the PC surface enhances the scattering cross-section of the nano-objects up to nearly 300-fold [16]. In addition, by illuminating the PC at the combination of wavelength and incident angle that results in nearly 100% reflection intensity (as dictated by the PC forbidden band) the transmitted reference light that combines with the forward-scattered light before reaching the image sensor is significantly attenuated [17]. Importantly, the enhanced scattering intensity provided by the PC resonance in PRISM enables the instrument to achieve high scattered intensity while using a relatively inexpensive, compact, and low-power (100 mW) laser compared to the Watt-level lasers used in several reported iSCAT instruments [2, 18-21]. Further, as the PC surface has the capability for directing out-scattered photons following its dispersion characteristic, the emerging scattered light is efficiently collected into the numerical aperture of an inexpensive and wide field of view (FOV) air-spaced objective lens (0.75NA), thus removing the need for oil-immersion, high numerical aperture objectives used in conventional iSCAT instruments [22-26]. The synergistic combination of PC surfaces as a substrate for performing iSCAT microscopy offers a route toward the detection and quantification of biological analytes in complex media, particularly when a selective recognition molecule is immobilized on the PC surface that enables it to capture a specific target. For example, using PRISM with PC surfaces prepared with an immobilized aptamer that selectively binds with the spike protein of SARS-CoV-2 virus, we demonstrated direct, rapid, 1-step, room temperature detection of SARS-CoV-2 in human saliva with detection limits similar to laboratory-based Polymerase Chain Reaction (PCR) analysis [27]. There are many potential applications for interferometric microscopy in the contexts of molecular diagnostics (for example, using gold nanoparticles (AuNPs) as tags for surface-captured molecules), viral diagnostics, and as a technology platform for a variety of life science research applications. These factors provide motivation to explore the development of compact and inexpensive instruments that can operate effectively without an optical bench, while automating some of the manual operations, such as the need to continuously maintain accurate focusing, to make the technique more accessible by users without an optics background.

Interferometric scattering microscopy faces inherent challenges for implementation outside of an optics laboratory due to the stringent requirements on the imaging condition. The interferometric nature of iSCAT and PRISM results in a bright-fringe system, where a weak signal intensity change must be detected on an overwhelming background [9]. This characteristic makes scattering-based microscopes vulnerable to signal fluctuations caused by external factors such as mechanical vibration and laser power instability. Therefore, most scattering microscopes are placed on air-damped optical tables for vibration isolation. Meanwhile, as the scattered signal scales with the intensity of illumination, scattering microscopes are often equipped with Class 4 lasers in the blue part of the optical spectrum [23, 28]. An efficient collection of scattered photons from inherently small objects generally dictates the selection of an oil-immersion objective lens [28, 29]. For diagnostic and life science research applications, where the detection instrument may be situated in a clinical laboratory or a biology-oriented laboratory, the environment places a limit on the space, budget, and available laser safety infrastructure, which renders some of these components impractical. Similar goals have motivated the development of portable fluorescence microscopes [30, 31].

In this report, we describe the design, implementation, and characterization of a PRISM instrument that is intended for use in environments besides optical laboratories, in which the system will provide sufficient contrast for the detection of nanoparticles, virions, exosomes, and large protein molecules while operating on a normal tabletop where extraneous vibration noise sources are present. The system incorporates the advantages reported initially for PRISM (reduced laser power, air-spaced objective) while also integrating an inexpensive (but effective) vibration isolation system based upon an elastic band suspension. The newly-developed instrument described in this report also incorporates an autofocus system that further stabilizes image contrast for extended time periods, while enabling the sensor to be rapidly translated to gather images from different locations, which facilitates the collection of “tiled” images that cover extended fields of view. The autofocus function is implemented by incorporating an additional light path to the sample without introducing significant background light intensity. After describing and characterizing the performance of the vibration isolation and autofocus capability, we characterize the system’s performance for sensing nano-objects by measuring the contrast from AuNPs with diameters in the 10-40 nm range. To demonstrate the capability of the system for observing various biological analytes, we challenged the system to detect HIV virus, SARS-CoV-2 virus, exosomes, and ferritin protein. In this work, the viruses are non-infectious pseudoviruses that retain the size and outer protein configuration of the native virus, without the presence of the infectious viral genome. Overall, we demonstrate a more portable and cost-effective version of the PRISM instrument that does not sacrifice detection capability for several important applications.

## 2. Materials and Methods

### 2.1 Instrument implementation

The optical assembly of the instrument can be separated into two major parts, namely the imaging component and the autofocus module, as shown in Fig. 1a. The imaging component uses a temperature-controlled 100mW (Class 3) laser diode (Thorlabs) emitting at 633 nm as the light source, which provides illumination to the sample in the Köhler configuration. The emitted light is then collimated, phase-shifted, and reshaped before entering the illumination objective lens (50X, NA = 0.5, Olympus) with polarization along the direction of the gratings on the PC surface. While the PC is illuminated at the resonance condition, only less than 0.5% of the incident light is transmitted, which interferes with the enhanced scattered light collected by the imaging objective lens (50X, NA = 0.75, Zeiss) (Fig.1b), forming an interferometric signal at the imaging sensor (Grasshopper3, Teledyne FLIR). The measured field of view (FOV) using a test target is 155 × 185 um^2^ with full-pixel utilization (2448 × 2048) at a frame rate of 75Hz. The FOV is then reduced to 512 × 512 pixels for a higher frame rate (530 Hz) and faster real-time data processing. The autofocus module uses an infrared laser diode (Edmunds Optics) emitting at 850 nm and is reshaped into a circular profile by an iris. A neutral density filter is applied to attenuate the autofocus laser power so that minimal light reaches the sample, avoiding unwanted background signal increase. The incident infrared light is reflected by a knife-edge prism mirror and a short-pass dichroic mirror (cutoff wavelength = 805 nm) before entering the back aperture of the imaging objective lens. The total internal reflected light follows the same path as the incident light, except it is reflected by the other side of the knife-edge prism mirror and is focused by a cylindrical lens onto the autofocus camera (Blackfly S, Teledyne FLIR) for further processing. A detailed component list can be found in Table S1.

**Fig. 1.**
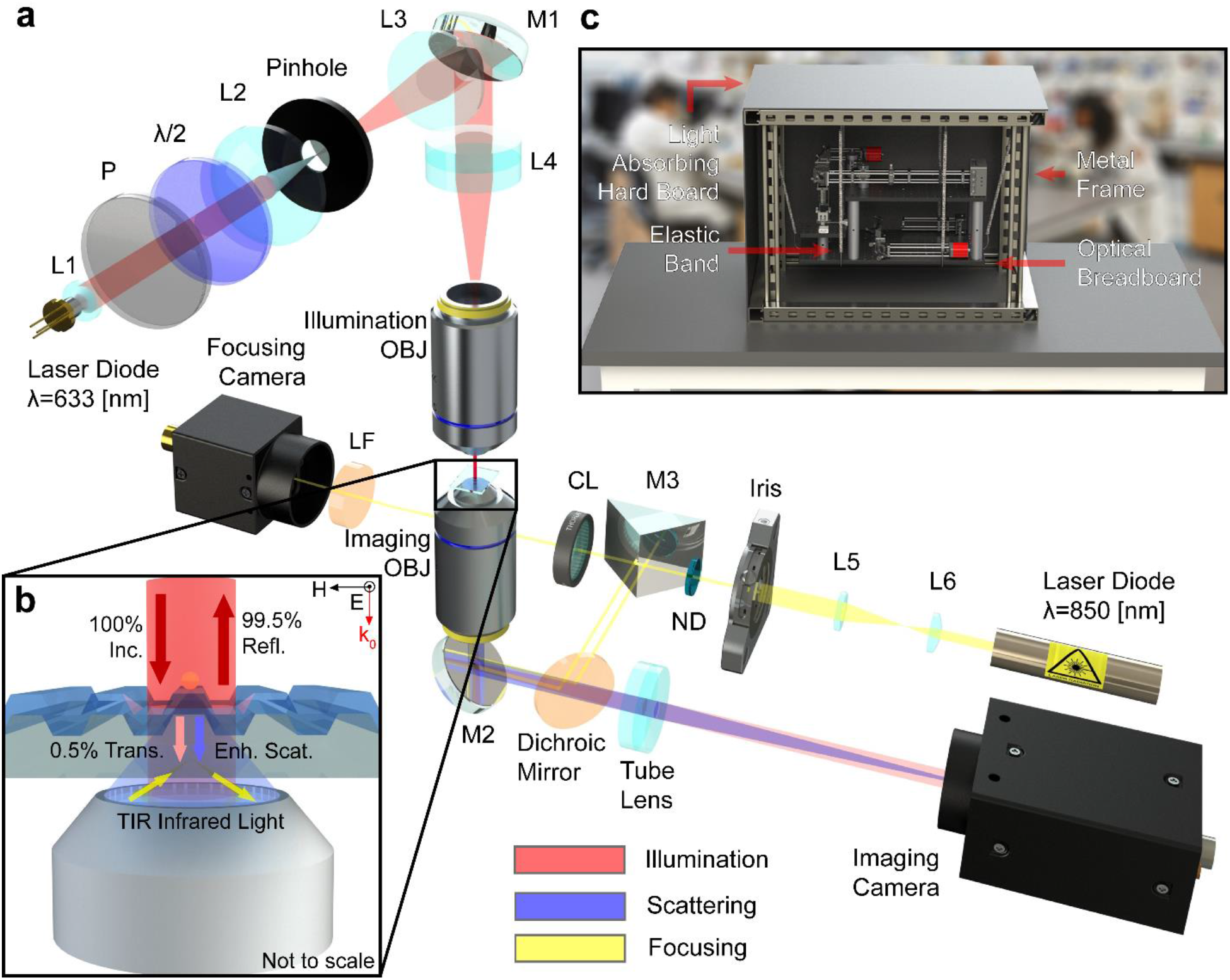
System schematics and working principles. (a) Schematic of the optical system. The 633 nm excitation light is colored in red, the scattered light from the sample is colored in blue, and the 850 nm light for the autofocus is colored in yellow. P polarizer, λ/2 half-wave plate, L1-6 spherical lens, M1-3 mirrors, OBJ objective lens, ND neutral density filter, CL cylindrical lens, LF laser line filter. (b) A magnified view near the sample. A TE-polarized collimated 633 nm laser beam illuminates the photonic crystal (PC) surface and excites the PC-guided resonance to enhance the scattered light. With the resonant condition satisfied, only 0.5% of light is transmitted and interferes with the enhanced scattering light, resulting in an interferometric signal on the imaging camera. The autofocus module also utilizes the same objective lens to form a total internal reflection of the 850 nm infrared light and senses the position of the sample. (c) A photo of the instrument in its housing, where the optical assembly is suspended by elastic bands to the metal frame and covered by light-absorbing material.

For the sake of vibration isolation and operator safety, the optical assembly is enclosed within a metal framework and is suspended by six elastic bands (Fig.1c). The elastic bands connect the optical breadboard to the girder and are stretched to ∼120% compared to their relaxed state. Light-absorbing hard boards cover the outer surface of the frame except for a small sliding window for sample loading. The enclosure blocks the ambient light from reaching the two cameras and prevents the laser light from escaping. The compact size of the instrument (15 × 27 × 18 in^3^) allows it to be placed on a wheeled table.

### 2.2 Photonic crystal fabrication

As reported in our previous work [16, 32, 33], the PCs are nanostructured surfaces comprised of linear gratings (period = 380nm, depth = 97 nm, duty cycle = 65%) etched into a glass substrate. The grating structure is patterned with holographic lithography on 8-inch diameter glass wafers, followed by reactive ion etching to form the grooves. An additional thin layer of TiO_2_ (thickness = 95nm), which acts as the guiding layer, is applied via sputter deposition (Fig.2a). The physical structure of the PC is designed such that low transmittance (<0.5%) is achieved under resonant wavelength when the PC surface is submerged in aqueous solution. Besides the attenuation to the transmitted light, PCs also exhibit surface-confined electromagnetic standing waves with an evanescent field extending into the liquid media, where the field intensity is comparatively stronger than the incident electric field (Fig. 2a). When nanoscopic scatterers travel within the enhanced electric field regions, they will experience an enhanced scattering cross-section; therefore, more photons are scattered and reach the imaging sensor. The resonant wavelength under this polarization is centered around 633 nm, overlapping with the laser illumination wavelength (Fig. 2b). The PCs used in this work are fabricated by a commercial vendor to our design specifications (Moxtek, Orem, UT) and provided as 10.16 × 12.7 mm^2^ chips. Each chip is subsequently cut with a dicing saw into four segments and mounted on microscope cover glass via ultra-violet curable adhesive for easier handling.

**Fig. 2.**
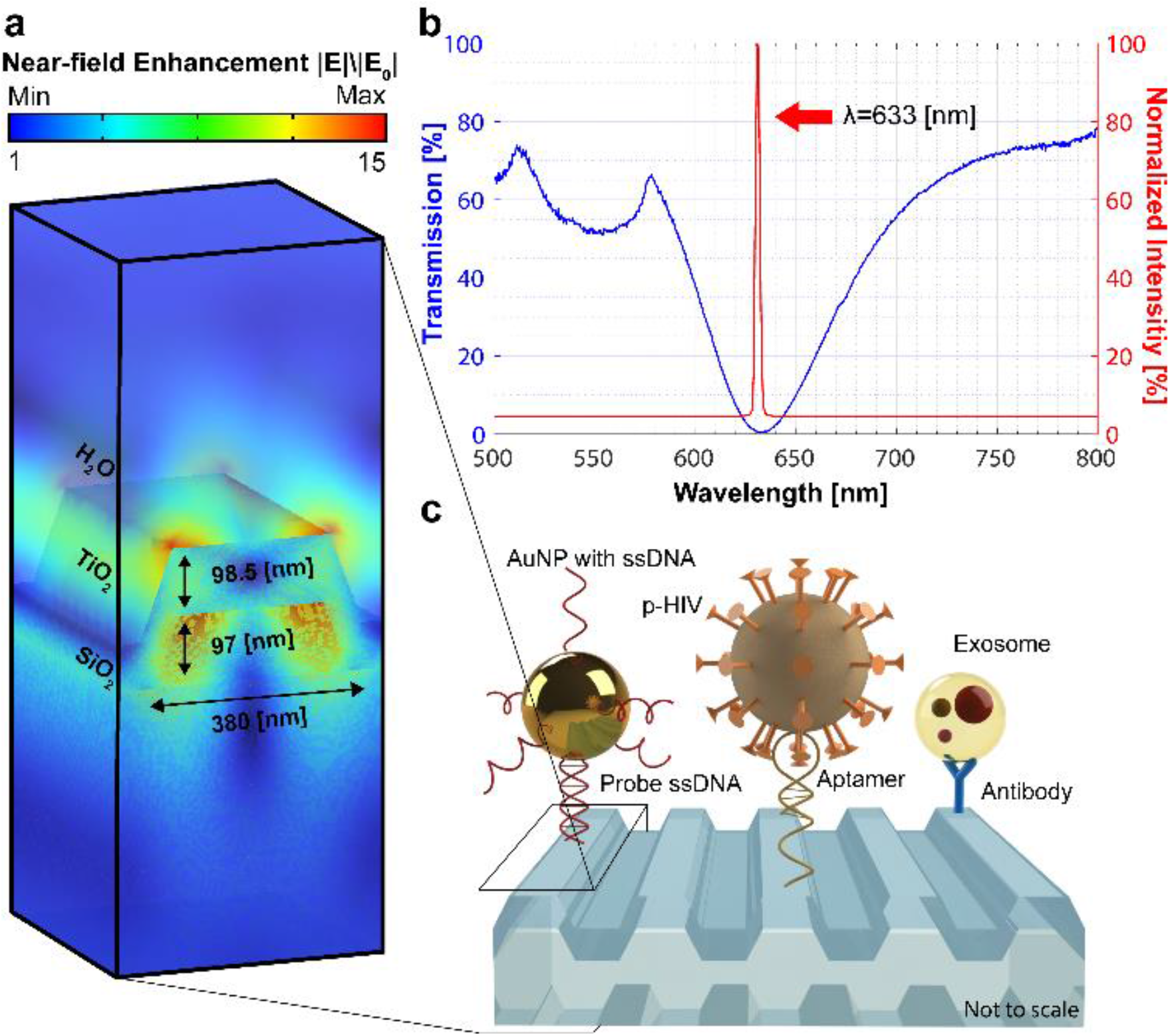
Photonic crystal properties and surface chemistry illustrations. (a) Normalized near-field electric field profile of one PC period under resonance. (b) The PC transmission and the normalized laser intensity spectrum. The blue curve represents the percentage of light intensity transmitted through a piece of PC under TE polarization. The lowest transmission occurs at λ = 633 nm, which corresponds to the resonant wavelength of the PC. The red curve represents the normalized laser intensity of a 100mW laser diode with a center wavelength of 633 nm. (c) Functionalized PC surface for capture and PRISM imaging of gold nanoparticles, HIV viruses, and exosomes.

### 2.3 Assay preparation

The PCs are cleaned by successively sonicating in acetone, isopropyl alcohol, and water for 1 minute each and dried via compressed nitrogen. They are further dried in an oven (60°C) before oxygen plasma treatment at 500 mTorr under 200 W power for 10 minutes (Pico Plasma System, Diener Electronic). After the PCs are plasma-cleaned, they are placed face down on a holding fixture within a glass jar with 50 uL of 3-Glycidoxypropyl trimethoxysilane (GPTMS) (Sigma Aldrich) at the bottom. The assembly is placed in a vacuum oven for 8 hours at 80°C to enable the GPTMS to enter the vapor phase, where the molecules can form covalent attachments with the exposed PC surface. After vapor-phase GPTMS coating, the PCs are removed from the oven and sonicated with methanol, toluene, and water for 1 minute each. For experiments performed to demonstrate the capture of exosomes with immobilized antibodies, instead of vapor-phase GPTMS, we performed liquid-phase silanization with 3-Aminopropyl triethoxysilane (APTES) (Sigma Aldrich), in which the PC was treated with 5% APTES in tetrahydrofuran solution for 1 hour, followed by a 10% dimethyl sulfide N,N’-Discuccinimidyl carbamate linker incubation for 45 minutes prior to antibody binding.

The five primary analytes in our experiments are AuNPs, HIV viruses, SARS-CoV-2 viruses, exosomes, and ferritin proteins. Each requires a slightly different immobilization procedure (Fig. 2c). For the AuNPs, the silanized PC surfaces are functionalized with DNA oligonucleotides (Integrated DNA Technologies) by incubating the PC in a 50 uM solution of amine-terminated ssDNA overnight. Complementary thiol-modified single-strand DNAs are conjugated to commercially available pre-activated AuNPs (10-40 nm, Cytodiagnostics) so that the nanoparticles will bind to the surface of the PCs. Pseudovirus capture follows a similar method; modified HD4 DNA aptamers (Integrated DNA Technologies) are fixed to the silanized PC surfaces by amine bonds, and these aptamers capture the HIV viruses via interactions with the GP120 protein on the outer shell of the virus [34]. The process for the capture of SARS-CoV-2 viruses resembles that of the HIV viruses, except AR-10 aptamers that selectively bind with the SARS-CoV-2 spike protein (Integrated DNA Technologies) are used [27]. The exosomes (retrieved from mouse plasma) are captured by CD147 antibodies that are functionalized on the PC surface. No additional linkage is needed for the ferritin molecules since the covalent bonds that directly form between the silanized PC surface and the free amine groups on the ferritin provide a stable bond to hold them in place. A more detailed surface chemistry description can be found in the supplementary information, and the sequences of oligonucleotides are documented in Table S2.

### 2.4 Data processing algorithms

The algorithms for image processing and autofocusing are written in MATLAB (MathWorks) and LabVIEW (National Instruments), respectively. The image processing algorithm is detailed in our previous report [16]. In brief, a rolling-window averaging method is applied to each frame to remove the constant background considering the dynamic measurements. The processed video shows the shot-noise-limited signals from the 3D movements of the nanoparticles (Video S1), rendering it ready for contrast signal recognition. A Laplacian-of-Gaussian filter is then applied to locate the local minima, followed by a 2D Gaussian function fitting for the point spread function. For each frame with identified particles, the algorithm calculates the contrast by dividing the signal by the pixel value obtained in the previous frame and then normalizing it. The contrast signal of a single detected particle in one frame is presented as one dot in the plot, as shown in Fig. 5b.

**Fig. 3.**
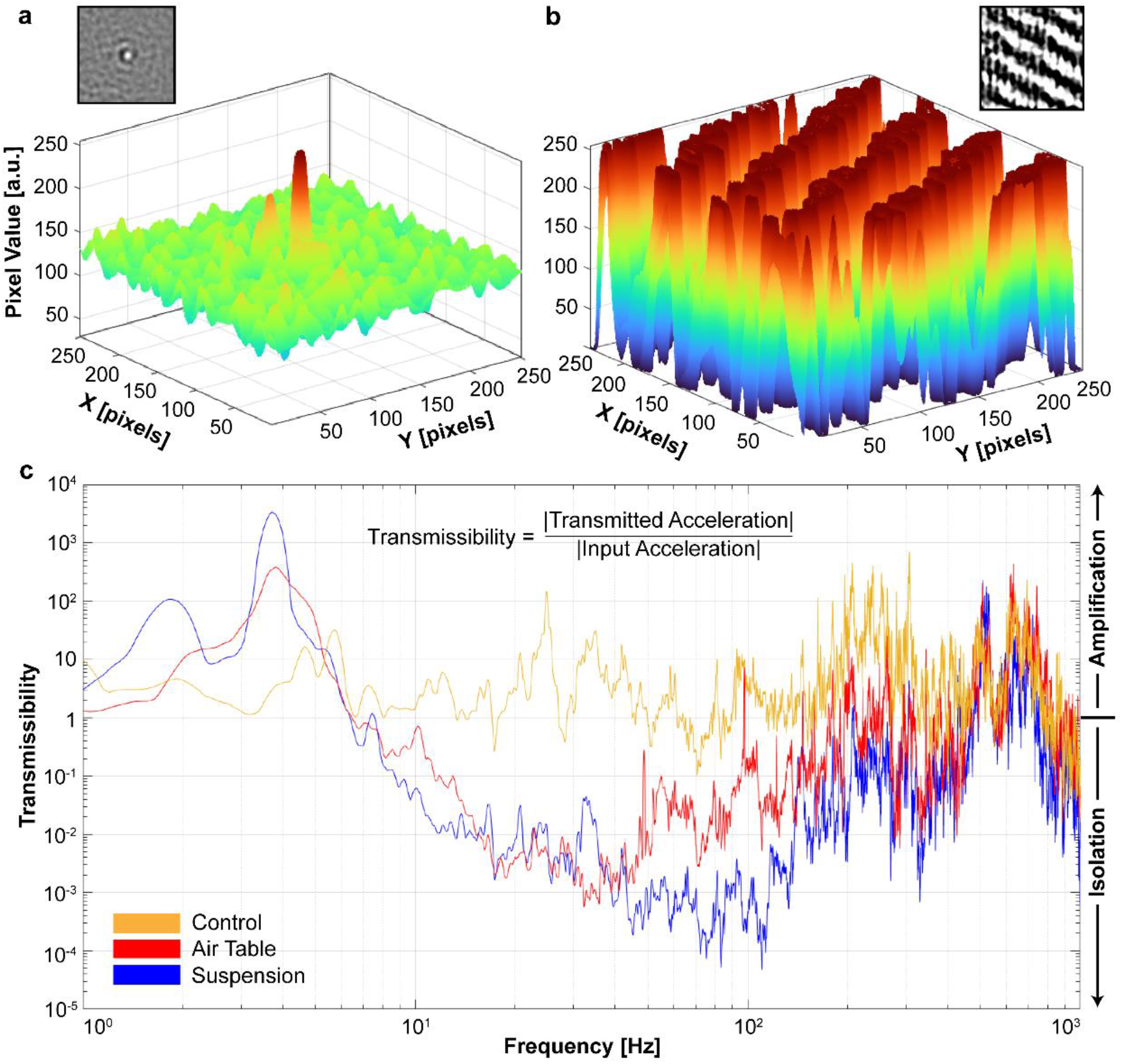
Vibration spectrum analysis. (a) Contour map of the differential image of an immobilized 40 nm AuNP with elastic bands suspending the imaging system. Inset: the plotted differential image. (b) Contour map of the differential image of an immobilized 40 nm AuNP without any vibration isolation method. Inset: the plotted differential image. (c) Transmissibility spectrum of an office desk (control, yellow), an air-damped optical table (red), and elastic band suspension (blue). A transmissibility above unity means the mechanical vibration is amplified, and a transmissibility below unity means the vibration is damped.

**Fig. 4.**
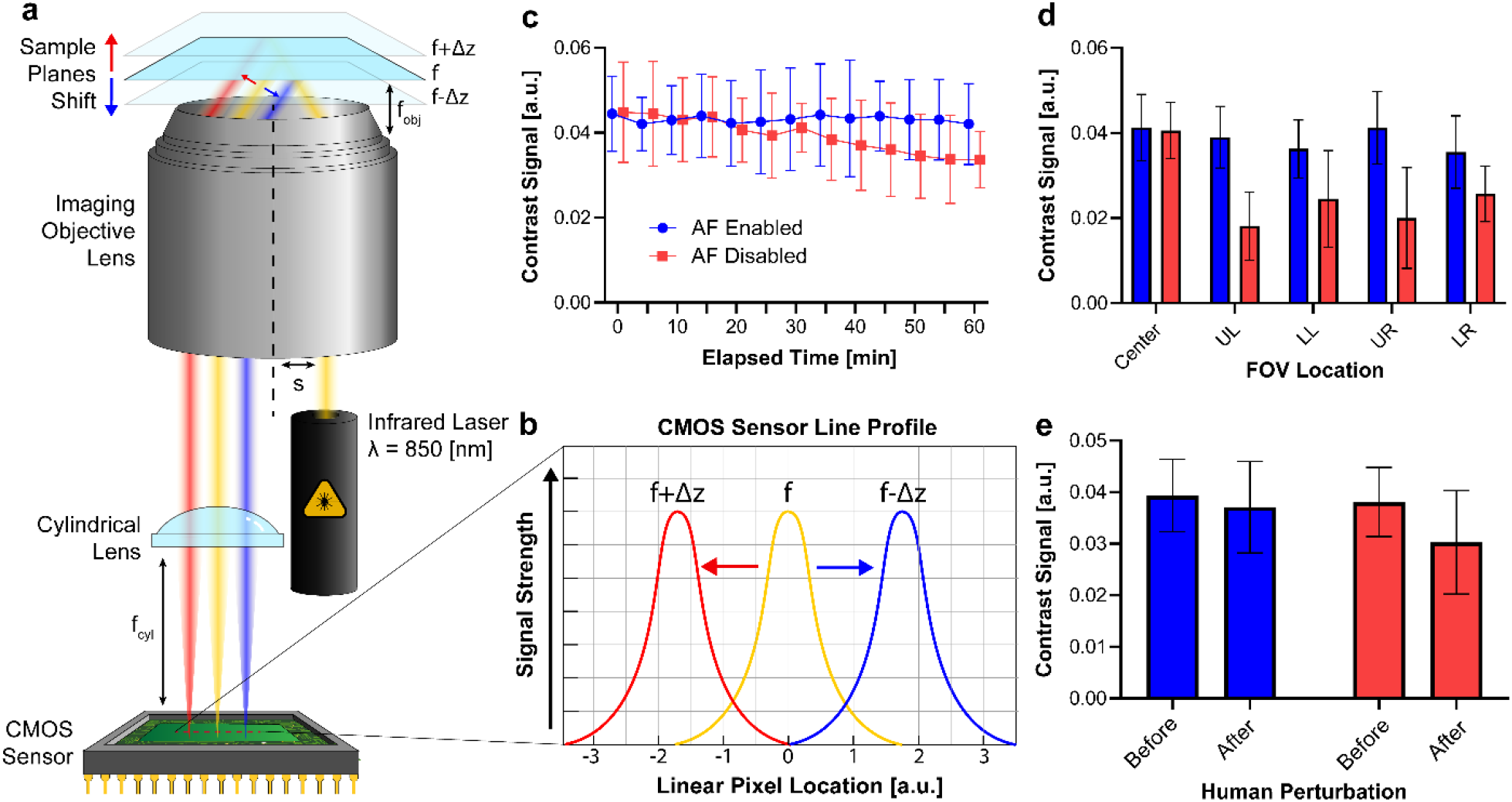
Automated focusing module illustration and characterization. (a) Illustration of the working principle of the autofocus module. The infrared beam emitted by a laser diode is totally reflected by the bottom surface of the microscope sample glass. As the sample drifts upward (represented by the red beam) or downward (blue beam), the Gaussian profile on the CMOS sensor will shift accordingly. (b) An exemplary line profile demonstrating how focal drift is translated to the shift of the peaks. (c) The change of contrast signals of captured 40 nm diameter gold nanoparticles over time with autofocus enabled (blue) and disabled (pink). (d) The change of contrast signals of captured 40 nm AuNPs over different locations across the PC surface when initially focused on the center of the sample. UL upper left, LL lower left, UR upper right, LR lower right. (e) The change of contrast signals after an intentional collision with the instrument.

**Fig. 5.**
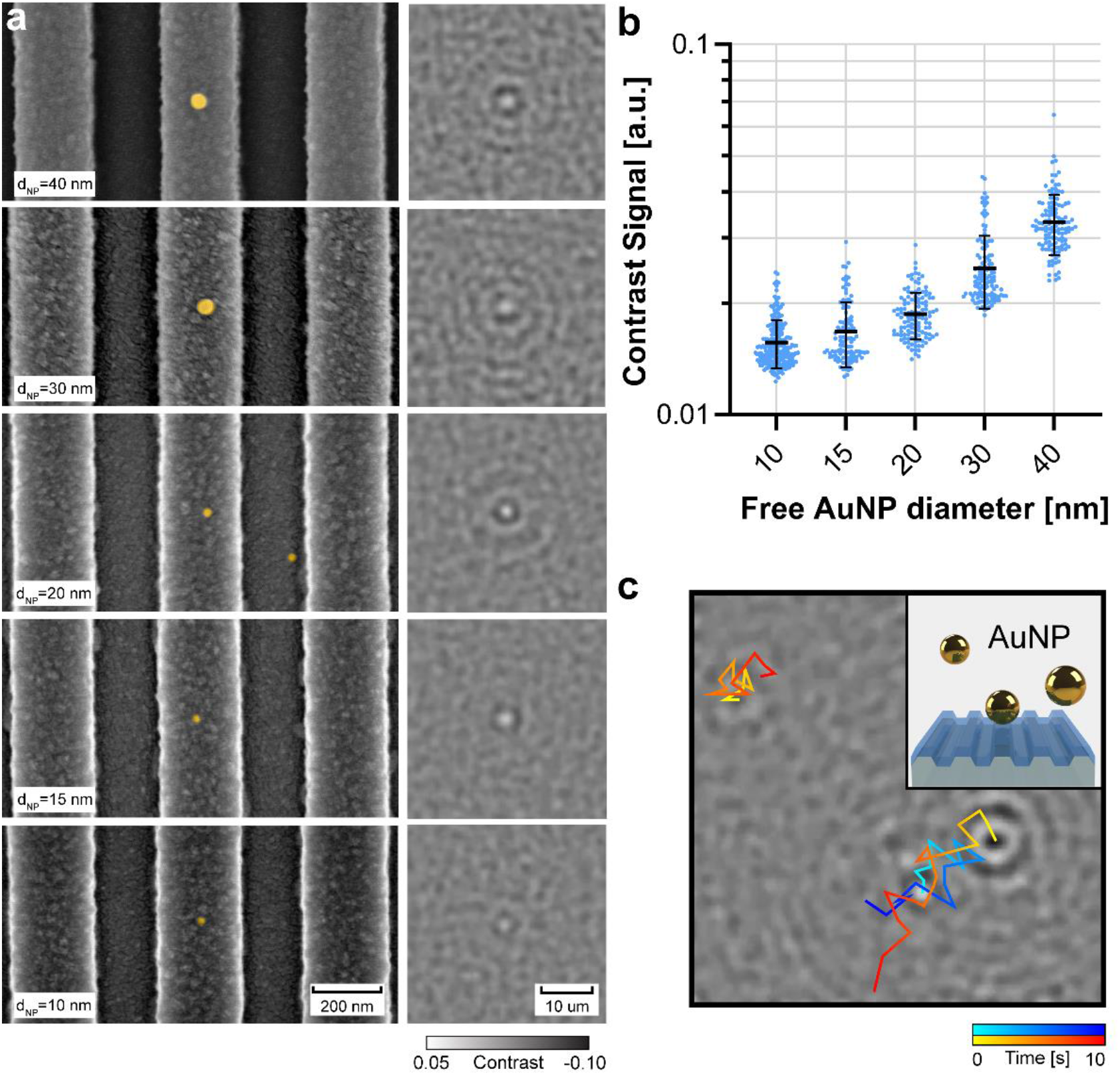
System characterization using free non-captured AuNPs. (a) False-colored SEM images of PC surfaces with AuNPs of various sizes ranging from 10 nm to 40 nm and their corresponding processed interferometric images. (b) Contrast signals of free-floating AuNPs of various sizes. Each dot represents the contrast of a detected AuNP in one frame. (c) Motion trajectory of three individual 40 nm AuNPs in a ten-second interval. Inset: An illustration of free-floating AuNPs with different distances to the PC surface.

A closed-loop system was built between the motorized stage and the focusing camera to implement the autofocus algorithm. The user sets the baseline and the tolerance for the peak position of the gaussian function on the camera line profile. Once a major focal dislocation occurs and the peak position falls out of the tolerance range, the algorithm will control the motorized stage to move responsively. A user-friendly interface is provided using the virtual instrument interface of LabVIEW, where the user can visualize and modify the CMOS camera line profile, set the parameters for autofocus, and manually control the motorized stage. A block diagram indicating the logic of the autofocus algorithm can be found in Fig. S1.

## 3. Results and discussion

### 3.1 Principle of imaging strategy

The imaging principle of the system resembles that of PRISM, and is fully described in our previous publication [16]. In short, the microscope is based on transmission iSCAT, where the transmitted light interferes with scattered light and forms interferometric signals at the imaging plane. Using PCs as sample substrates and illuminating them under resonant conditions, the system achieves three benefits: 1. Propagating leaky modes create evanescent fields that enhance the scattering cross-section. 2. Attenuated transmitted reference light increases the contrast detected by the camera. 3. Resonance-guided angular scattering increases the number of scattered photons collected by the imaging objective lens. The benefits brought by the PCs allow us to use air-spaced low-NA objective lenses, low-performance cameras, and Class 3 red laser sources for imaging purposes. The contrast signal calculated by the algorithm is positively correlated with the volume of the particle of interest and can be expressed as:

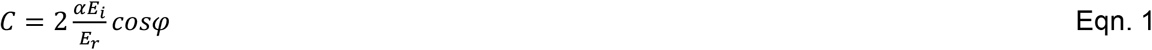

where *α* = 3*ϵ*_*m*_*V*(*ϵ*_*p*_ − *ϵ*_*m*_)/(*ϵ*_*p*_ + *ϵ*_*m*_) is the complex particle polarizability [35], *E*_*i*_ is the incident electric field magnitude, *E*_*r*_ is the referencing electric field magnitude, *φ* is the phase difference between the two fields, *V* is the particle volume, and *ϵ*_*p*_ and *ϵ*_*m*_ are the permittivities of the particle and the embedding medium. However, for our instrument, the linear relationship between the contrast and volume of our system is slightly compromised by the presence of localized surface electric field hot spots on the PCs due to the random capture sites. The enhancement of the scattering cross section also increases as the particle size shrinks [16], which yields another level of nonlinearity. Therefore, we do not seek to estimate the mass of the detected nano-objects through the magnitude of the image contrast. Rather, the system detects the presence of the analytes and provides an insight into their relative sizes, making this instrument ideal for the characterization of specific captured objects and their qualitative size ranges.

### 3.2 Vibration isolation characterization

One major obstacle to scattering microscopes for clinical applications is the lack of a lightweight but effective vibration isolation method. An air-damped optical table is the most common choice for optical instruments; however, its lack of portability and large footprint make it unsuitable for many environments such as clinical diagnostic laboratories and biology laboratories. Novel vibration isolation methods, such as the use of magnetorheological elastomers as active dampers [36], require careful design and meticulous care for long-term stability. Inspired by vibration isolation methods used for scanning tunneling microscopes [37, 38], we sought to utilize the classical approach of using elastic bands as passive vibration dampers to meet our requirements. With the imaging system suspended by elastic bands, the noise caused by mechanical vibrations is insignificant and covered by photon shot noise and signals from out-of-plane nanoparticles (Fig. 3a). The low background noise level in the differential image provides a large signal-to-noise ratio and facilitates the identification of the nano-objects. However, when the instrument is placed directly on an office table, the mechanical vibration saturates the pixel value range and overwhelms the AuNP signal on the differential image (Fig. 3b), making it impossible to identify the nanoparticles. To quantitatively compare the vibration isolation methods, we measured the vibration of the optical assembly when it was sitting on an office desk (control), suspended by elastic bands, and placed on an air-damped optical table (CleanTop II, TMC) with a high-precision accelerometer (731A, Wilcoxon). We also measured the input acceleration (vibration) by placing the accelerometer near the legs of the office desk and the optical table, respectively. The temporal vibration data is Fourier transformed into the frequency domain and then divided by the input acceleration to calculate the transmissibility. The three datasets are triangularly smoothed to display the amplitude at different frequency ranges, namely 1-10 Hz (low), 10-100 Hz (medium), and 100-1000 Hz (high) (Fig. 3c). In comparison to the control group, the elastic band and the air table exhibit a relatively higher damping efficiency over the entire vibration spectrum (Fig. S2a), except for extremely low frequencies less than 6 Hz. To produce a close resemblance to environments where human movements may cause additional mechanical vibrations, we stepped on the ground at a constant pace of 60 steps/min, one meter away from the instrument, simulating people walking in the immediate area. The experiment was repeated ten times, and the averaged result shows that the elastic suspension method has stronger resistance to footsteps in all frequency ranges, except for small disadvantages (0.7 dBV in magnitude) in the mid-range compared to the air table. (Fig. S2b).

### 3.3 Automized focusing characterization

A handful of commercially available products support autofocus functions on modern microscopes, for example, the Definite Focus from Zeiss and the Perfect Focus System from Nikon. However, these products are either only compatible with their branded microscopes or too expensive to be considered in a point-of-use instrument application. The autofocus module we integrated with our scattering microscope is based on similar principles as the commercial products; the eccentric light beam that enters the back aperture of the objective lens reaches the sample at an oblique angle. This incident angle is larger than the critical angle, and all light is reflected to the imaging objective lens. A cylindrical lens then focuses the reflected beam to form a line pattern on the CMOS imaging sensor; meanwhile, a 1 × 3072 pixels linear profile is taken perpendicular to the line patterns for analysis. The reflected line pattern exhibits a quasi-Gaussian distribution, where its peak is used to locate the central position. Once a sample drift occurs, denoted as ± Δ*z*, the peak position of the Gaussian distribution will shift accordingly (Fig. 4a). Consequently, the number of shifted pixels can be used to determine the magnitude of the focal drift and in what direction the sample stage must move to correct the focus. The distance that the line pattern shifts at the image plane can be expressed as [39]:

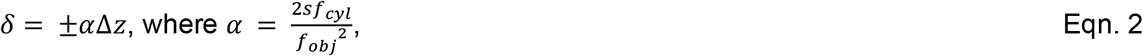

Where *s* is the offset distance to the optical axis, *f*_*cyl*_ is the focal length of the cylindrical lens, and *f*_*obj*_ is the focal length of the objective lens. The calculated *α* for our system is ∼2000, which means that the focal shift is magnified 2000 times and translated onto the image sensor. With a 2.4 × 2.4 um^2^ pixel size, the calculated resolution for the focal drift is around 1.2 nm. However, this limit is compromised by the sensor’s dark current, photon shot noise, and the piezoelectric sample stage due to its large step size (∼1 um).

We investigated the performance of the autofocus module using captured 40 nm diameter AuNPs as targets and conducted tests on the temporal and spatial contrast signal change. Only one AuNP was present in the FOV, and a 10-second video was recorded every 5 minutes in a 60-minute interval for the temporal test. The result with autofocus disabled showed a consistent decrease in the contrast signal over time at an average rate of 1.86 × 10^−4^ per minute, whereas the value with autofocus enabled is stable (Fig. 4c). To test the spatial dependence of the contrast signal with or without autofocus, we first manually focused on the center of the PC sample, and then move the FOV to the four corners. Without the autofocus being activated, all four corners showed degraded contrast signals at an average of 0.0159. In comparison, the autofocus-aided trials had a well-maintained contrast level (Fig. 4d). Finally, we dropped a 1 kg weight from 0.5 m above the table that held the imaging system to test the self-correction of the autofocus module after an artificial collision. While the autofocus module was able to recover the focal position, there is still a minor degradation of the contrast signal (Fig. 4e).

### 3.4 System calibration with AuNPs

To characterize the size differentiation capability of the system, we detected AuNPs with diameters varying from 10 to 40 nm. A droplet of liquid (∼ 5uL) that contains 100× diluted AuNPs (∼1.5×10^9^ particles/ml) is uniformly spread across the entire PC surface by clipping a cover glass on the top. Because of their plasmonic nature, AuNPs generally possess a larger scattering cross section compared to biological substances of the same size [40], resulting in a higher contrast signal on the image. Scanning electron microscope images of the PC’s groove structures with false-colored AuNPs are shown in the first column of Fig. 5a. The second column corresponds to their processed images obtained with our scattering microscope. It can be observed that as the size gets smaller, the central lobe becomes fainter, as represented in a contrast signal plot against various particle sizes (Fig. 5b). Importantly, the AuNPs that we used as size calibration targets are free to move in all directions (Video. S1); therefore, the large standard deviation is a result of particles moving into and out of the focal plane and their relative distance to the PC surface. The trajectories of three uncaptured 40 nm AuNPs are traced in a 10-second video to indicate their free-floating nature (Fig. 5c). However, Fig. 5b still proves that our imaging system can analytically compare the size of the nano-objects and that the diameter of the smallest free AuNPs that it can resolve is 10 nm.

### 3.5 Demonstration with various biological samples

In addition to AuNPs, we tested the imaging capability of our system with biological samples of clinical relevance, namely HIV viruses, SARS-CoV-2 viruses, ferritin proteins, and exosomes. HIV and SARS-CoV-2 viruses are common analytes in medical labs because they can be found in the patient’s bodily fluid and are crucial to early-stage HIV and COVID-19 disease diagnostics. Ferritin is useful for this study because it has significant roles in preserving iron in vertebrates and it is a relatively large protein (∼440 kDa) with a core consisting of up to 4000 iron atoms [41, 42]. Other studies based on iSCAT have targeted ferritin as a large protein and it was found to have a contrast signal commensurate with its mass [12, 43]. Exosomes are clinically relevant because they could be potential biomarkers for cancer diseases [44]. As a subset of extracellular vesicles, exosomes are liposomal nanovesicles secreted by mammalian cells that range from 30-200 nm in diameter [45]. They are essential parts of intercell communications, carrying important substances, such as transmembrane proteins, mRNA, and miRNA, from the original cell [46]. Several aptamer-based biosensors have demonstrated the detection capability of exosomes, including fluorescence [47], electrochemical [48], and surface plasmon resonance methods [49]. However, limited scientific literature reports the digital imaging of exosomes with variants of iSCAT up to date [14].

All four types of particles were immobilized to the PC surface with methods described in section 2.3; consequently, their movements were insignificant compared to free particles. This greatly aided the tracking of small particles at low concentrations because it would not require continuous FOV shift to chase the particle’s movement. Among the four substances, ferritin has the lowest contrast signal, which is expected considering the molecular weight of these proteins. The comparatively small standard deviation of the ferritin contrast signals can be explained by the fact that they were directly bonded to the PC surface and minimal vertical movements occurred. The exosomes extracted from mouse plasma have an average diameter of ∼100 nm (Fig. S2). Both HIV and SARS-CoV-2 pseudoviruses are based on lentivirus cores, giving them approximately the same diameters (∼150 nm) [27]. Contrary to their similar dimensions, the contrast signal of SARS-CoV-2 viruses is observed to be higher than that of HIV viruses (Fig. 6a). To explain this strange phenomenon, we performed surface plasmon resonance (SPR) binding analysis on AR10 aptamers (selected against the entire SARS-CoV-2 viral particle) [50] and RBD-1C (selected against spike RBD) [51] (Fig. S4). The SPR results suggested that AR10 did not interacts with SARS-CoV-2 spike proteins, and since the spike proteins are the only protruding proteins on the virion’s outer surface, we concluded that AR10 aptamer directly interacts with some moiety on the viral membrane. This brought the virions closer to the PC surface and thus the contrast signal is increased due to localized surface-enhanced electric field regions. We also investigated the enhancement of the scattered light by comparing the contrast signal of free and captured particles. Stronger signals are detected for immobilized particles because they are brought into closer proximity with the PC surface and experience a stronger field intensity. The contrast signal increases by 5.30 × 10^−3^ on average for 40 nm AuNP (Fig. 6b) and 3.75 × 10^−3^ for HIV viruses (Fig. 6c), proving that the surface-enhanced electric field areas created by the leaky guiding modes indeed improve scattering cross-sections.

**Fig. 6.**
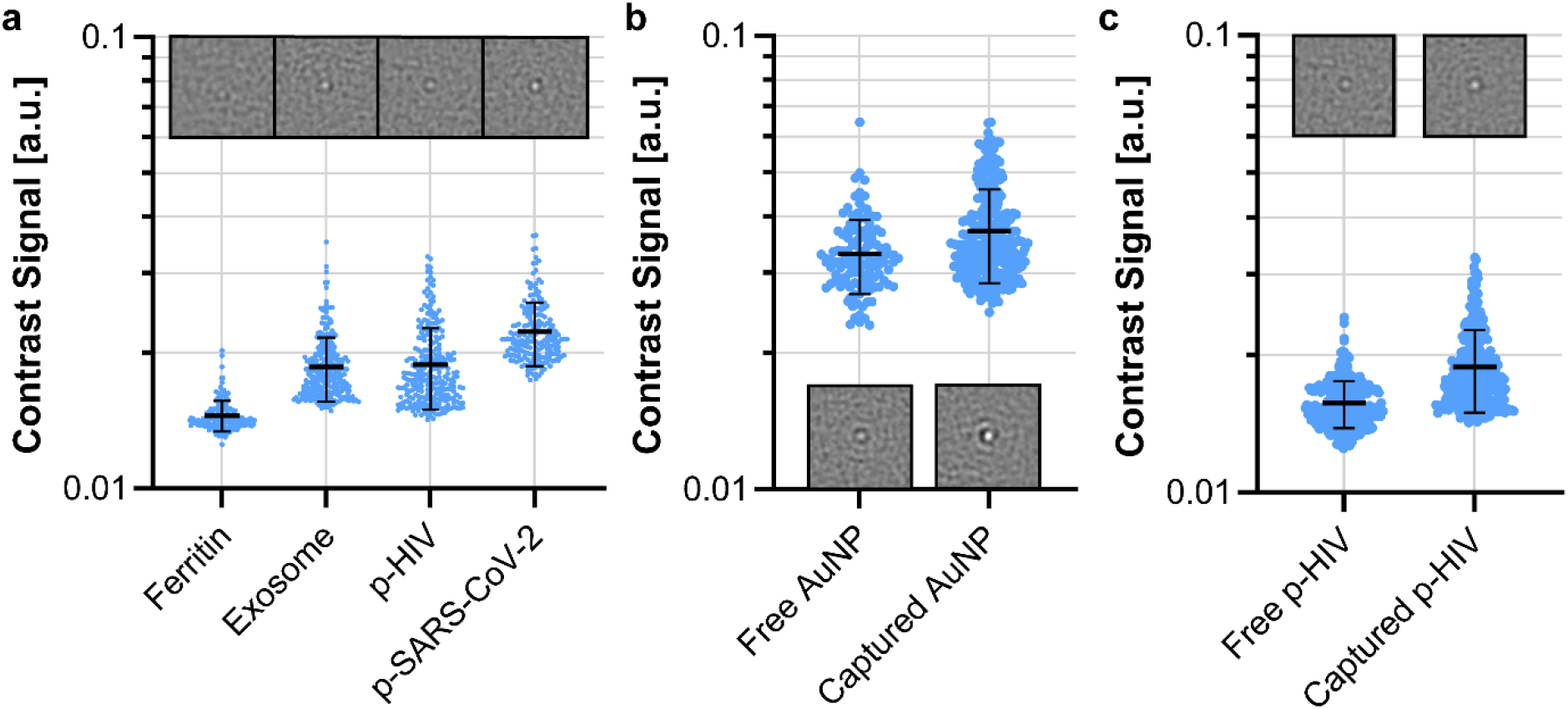
Experiments demonstrating the imaging capability. (a) Contrast signal distribution of various immobilized biological nanoparticles with their interferometric images as insets. Ferritin molecule(440 kDa), Exosome (∼100 nm), HIV virus (∼150 nm), SARS-CoV-2 virus (∼150 nm). (b) Contrast signal comparison between free and captured 40 nm AuNPs. (c) Contrast signal comparison between free and captured HIV viruses.

## 4. Conclusions

In conclusion, a scattering microscope designed for point-of-use environments outside an optics laboratory was constructed, and its performance was tested. The autofocus module maintains a stable signal strength over time and spatial position, establishing a foundation for large-area scanning applications. The vibration isolation method using an elastic band suspension shows an overall better resistance to mechanical vibrations than an optical air table, creating new possibilities for portable and cost-effective microscope systems that are vibration sensitive. The imaging system is proven capable of differentiating a range of different-sized AuNPs and imaging clinically relevant substances such as proteins, exosomes, and virions at digital resolution. All the above experiments justify its performance in a point-of-use environment and ability to detect potentially relevant biological analytes. Future improvement can be made by honing the detection algorithm for live particle recognition and incorporating z-scanning to extract phase information for particle categorization [52]. Efforts can also be made to minimize the system to a size comparable to a desktop computer for applications requiring additional portability. Expanding the application of iSCAT to settings such as clinical diagnostic facilities and biology laboratories opens the possibility for advanced point-of-care diagnostic systems, and this device is a promising step toward achieving that goal.

## Acknowledgment

The authors thank Dr. Lijun Rong and Dr. Laura Cooper for providing the pseudoviruses, and Prof. Marni Boppart and Dr. Ray Spradlin for providing the exosomes. The authors are grateful to Prof. Joseph Lyding for important suggestions for implementing the elastic suspension and for sharing the accelerometer used for vibration spectrum measurements. The authors are grateful for financial support from NIH (R01AI159454) and CSL Behring. JT acknowledges support from an NSF Graduate Fellowship.

