## Supplementary material for "A Photonic Resonator Interferometric Scattering Microscope for Label-free Detection of Nanometer-Scale Objects with Digital Precision in Point-of-Use Environments": Supplementary material.pdf

### Supplementary Information

| Component Name | Quantity | Unit Price ($) | ITEM# | Total Price ($) |
|---|---|---|---|---|
| Aluminum Breadboard, 18" x 24" x 1/2", 1/4"-20 Taps | 1 | 408.85 | MB1824 | 408.85 |
| Aluminum Breadboard 12" x 24" x 1/2", 1/4"-20 Taps | 1 | 288.15 | MB1224 | 288.15 |
| Ø1.5" Mounting Post, 1/4"-20 Taps, L = 14" | 4 | 101.82 | P14 | 407.28 |
| TE-Cooled Mount for Ø5.6 mm Laser Diodes with F or G Pin Codes, 1/4"-20 Taps | 1 | 706.83 | LDM56F | 706.83 |
| 633 nm, 100 mW, Ø5.6 mm, G Pin Code, Laser Diode | 1 | 141.04 | HL63163DG | 141.04 |
| K-Cube™ Laser Diode Driver | 1 | 909.29 | KLD101 | 909.29 |
| ±15 V/5 V Power Supply Unit with Mini-DIN Connectors for up to Two K- or T-Cubes | 1 | 119.78 | TPS002 | 119.78 |
| T-Cube TEC Controller, 4W | 1 | 815.24 | TTC001 | 815.24 |
| Mounted Ø7.200mm, f=5.5mm, NA=0.60 D-ZK3 Asphere,-A Coated | 1 | 132.93 | C105TMD-A | 132.93 |
| Kinematic 30 mm-Cage-Compatible Mount for Ø1" Optic | 2 | 101.77 | KC1 | 203.54 |
| 30.0mm Cage Plate to M9 Lens Adapter | 1 | 40.04 | CP1M09 | 40.04 |
| Right-Angle Kinematic Mirror Mount with Smooth Cage Rod Bores, 30 mm Cage System | 6 | 147.29 | KCB1C | 883.74 |
| Ø1" Protected Silver Mirror, 10 Pack | 1 | 462.93 | PF10-03-P01-10 | 462.93 |
| Cage Assembly Rod, 8" Long, Ø6 mm, 4 Pack | 2 | 45.79 | ER8-P4 | 91.58 |
| Cage Assembly Rod, 6" Long, Ø6 mm, 4 Pack | 3 | 33.86 | ER6-P4 | 101.58 |
| Cage Assembly Rod, 4" Long, Ø6 mm, 4 Pack | 3 | 27.79 | ER3-P4 | 83.37 |
| Cage Assembly Rod, 2" Long, Ø6 mm, 4 Pack | 4 | 23.88 | ER2-P4 | 95.52 |
| Cage Assembly Rod, 1" Long, Ø6 mm, 4 Pack | 3 | 19.77 | ER1-P4 | 59.31 |
| Rod Adapter for Ø6 mm ER Rods, L = 0.27", 4 Pack | 2 | 60 | ERSCB-P4 | 120 |
| SM1-Threaded 30 mm Cage Plate, 0.35" Thick, 2 Retaining Rings, 8-32 Tap | 6 | 18.18 | CP33 | 109.08 |
| Ø1" N-BK7 Plano-Convex Lens, SM1-Threaded Mount, f = 30 mm, ARC: 350-700 nm | 1 | 50.27 | LA1805-A-ML | 50.27 |
| Ø1" N-BK7 Plano-Convex Lens, SM1-Threaded Mount, f = 150 mm, ARC: 350-700 nm | 1 | 47.97 | LA1433-A-ML | 47.97 |
| 30 mm Cage System, XY Translating Lens Mount for Ø1" Optics | 2 | 191.88 | CXY1A | 383.76 |
| 30 mm Cage System, XY Translating Mount for Ø1" Optics with Quick Release Plate | 1 | 255.11 | CXY1QA | 255.11 |
| Ø1" Mounted Pinhole, 50 ± 3 µm Pinhole Diameter, Stainless Steel | 1 | 73.41 | P50K | 73.41 |

| | | | | |
|---|---|---|---|---|
| f=100 mm, Ø1" Achromatic Doublet, SM1-Threaded Mount, ARC: 400-700 nm | 1 | 108.7 | AC254-100-A-ML | 108.7 |
| Ø1" N-BK7 Plano-Convex Lens, SM1-Threaded Mount, f = 100 mm, ARC: 350-700 nm | 1 | 48.53 | LA1509-A-ML | 48.53 |
| 30 mm to 60 mm Cage Plate Adapter, 8-32 Tap | 2 | 43.26 | LCP33 | 86.52 |
| Adapter with External RMS Threads and Internal SM1 Threads | 2 | 24.43 | SM1A4 | 48.86 |
| ORIC 20 mm Monolithic XY Stage with Piezoelectric Inertia Drive, Imperial | 1 | 1568.25 | PD1D | 1568.25 |
| ORIC 20 mm Linear Stage with Piezoelectric Inertia Drive, Imperial | 1 | 728.47 | PD1 | 728.47 |
| Right-Angle Bracket Adapter for 20 mm Piezo Inertia Stage, Imperial | 1 | 88.15 | PD1Z | 88.15 |
| Four-Channel K-Cube Piezo Inertia Motor Controller | 1 | 1127.6 | KIM101 | 1127.6 |
| 15 V, 2.66 A Power Supply Unit with 3.5 mm Jack Connector for One K- or T-Cube | 1 | 37.66 | KPS201 | 37.66 |
| f=250 mm, Ø1" Achromatic Doublet, SM1-Threaded Mount, ARC: 400-700 nm | 1 | 108.7 | AC254-250-A-ML | 108.7 |
| C-Mount-Threaded 30 mm Cage Plate, 0.35" Thick, 8-32 Tap | 2 | 33.6 | CP13 | 67.2 |
| FLIR Grasshopper 3 USB3, 2.3 MP, 163 FPS, Sony IMX174, Mono | 1 | 1179 | GS3-U3-23S6M-C | 1179 |
| 50X ZEISS EC Epiplan Objective | 1 | 1118 | 13-826 | 1118 |
| 50X OLYMPUS LMPLFLN Objective | 1 | 2717.61 | LMPLFLN | 2717.61 |
| 25mW, 850nm Alignment Laser Diode | 1 | 199 | 19-454 | 199 |
| 5V DC Power Supply | 1 | 56 | 83-855 | 56 |
| Ø1" Unthreaded Adapter for Ø12 mm Cylindrical Components | 1 | 24.96 | AD12NT | 24.96 |
| SM05-Threaded 30 mm Cage Plate, 0.35" Thick, Two Retaining Rings, 8-32 Tap | 1 | 19.29 | CP32 | 19.29 |
| Ø1/2" N-BK7 Plano-Convex Lens, SM05-Threaded Mount, f = 25 mm, ARC: 650-1050 nm | 1 | 48.53 | LA1560-B-ML | 48.53 |
| Ø1" N-BK7 Plano-Convex Lens, SM1-Threaded Mount, f = 50 mm, ARC: 650-1050 nm | 1 | 49.92 | LA1131-B-ML | 49.92 |
| 30 mm Cage System Iris Diaphragm (Ø0.8 - Ø20 mm) | 1 | 97.38 | CP20D | 97.38 |
| Kinematic Mirror Mount for Ø1" Optics | 2 | 39.86 | KM100 | 79.72 |
| Right-Angle Kinematic Mount for Ø1" Optics, 30 mm Cage Compatible | 1 | 168.55 | C45P | 168.55 |
| Ø1" Shortpass Dichroic Mirror, 805 nm Cutoff | 1 | 289.5 | DMSP805 | 289.5 |

| | | | | |
|---|---|---|---|---|
| Ø2" Manual Rotation Stage | 1 | 104 | RP01 | 104 |
| Knife-Edge Right-Angle Prism Prot. Silver Mirror, 450 nm-20 µm, L = 25 mm | 1 | 136.43 | MRAK25-P01 | 136.43 |
| Platform Mount for 1" or 25.0 mm Beamsplitters and Right-Angle Prisms, Imperial | 1 | 51.02 | BSH1 | 51.02 |
| Cage Rotation Mount for Ø1" Optics, SM1 Threaded, 8-32 Tap | 3 | 89.57 | CRM1T | 268.71 |
| f = 250.0 mm, Ø1", N-BK7 Mounted Plano-Convex Round Cyl Lens, ARC: 650 - 1050 nm | 1 | 122.02 | LJ1267RM-B | 122.02 |
| Ø1" Linear Polarizer with N-BK7 Windows, 400-700 nm | 1 | 100.93 | LPVISE100-A | 100.93 |
| Ø1/2" Zero-Order Half-Wave Plate, Ø1" Mount, 633 nm | 1 | 473.62 | WPH05M-633 | 473.62 |
| FLIR USB 3.1 Blackfly® S, Monochrome Camera | 1 | 495 | BFS-U3-63S4M-C | 495 |
| SM05 Lens Tube, 0.50" Thread Depth | 1 | 14.52 | SM05L05 | 14.52 |
| Ø25 mm Absorptive Neutral Density Filter, ARC: 650-1050 nm, SM1-Threaded Mount, OD: 3.0 | 1 | 81 | NE30A-B | 81 |
| Ø1.85" Studded Pedestal Base Adapter, 1/4"-20 Thread | 4 | 14.08 | PB4 | 56.32 |
| Clamping Fork for Ø1.5" Pedestal Post or Post Pedestal Base Adapter, 1.25" Counterbored Slo | 4 | 15.38 | PF125B | 61.52 |
| Clamping Fork, 1.24" Counterbored Slot, Universal, 5 Pack | 1 | 43.72 | CF125-P5 | 43.72 |
| Tapped Mounting Base, 2" x 4" x 3/8" | 1 | 28.27 | BA2L | 28.27 |
| Ø1" Pedestal Pillar Post, 8-32 Taps, L = 4" | 3 | 35.22 | RS4P8E | 105.66 |
| Right-Angle Ø1" to Ø1/2" Post Clamp | 3 | 29.12 | RA90RS | 87.36 |
| 1/2" Optical Post, SS, 8-32 Setscrew, 1/4"-20 Tap, L = 2", 5 Pack | 3 | 25.26 | TR2-P5 | 75.78 |
| Ø1/2" Optical Post, SS, 8-32 Setscrew, 1/4"-20 Tap, L = 6", 5 Pack | 1 | 34.64 | TR6-P5 | 34.64 |
| Ø1/2" Post Holder, Spring-Loaded Hex-Locking Thumbscrew, L = 3", 5 Pack | 1 | 43.65 | PH3-P5 | 43.65 |
| Ø1/2" Post Holder, Spring-Loaded Hex-Locking Thumbscrew, L=1.5", 5 Pack | 3 | 38.11 | PH1.5-p5 | 114.33 |
| Phoenix Support Systems 1-5/8" X 1-5/8" X 10' Galvanized Strut Channel | 5 | 34.48 | A12H1000PGR | 172.4 |
| 1/2" x 18" Black Bungee Cords (bundle of 10) - 12mm | 1 | 49.99 | NA | 49.99 |
| Black Posterboard, 20" x 30" (508 mm x 762 mm), 1/16" (1.6 mm) Thick, 5 Sheets | 2 | 51.31 | TB5 | 102.62 |
| Matte Black Aluminum Foil, 1' x 50' (305 mm x 15.2 m) x .002" (50 µm) Thick | 1 | 33.95 | BKF12 | 33.95 |
| High-Performance Black Masking Tape, 2" x 180' (50 mm x 55 m) Roll | 1 | 43.26 | T743-2.0 | 43.26 |

| | | | | |
|---|---|---|---|---|
| | | | | 19957.42 |

Table S1. Component list.

| Name | Sequence (5' to 3') |
|---|---|
| AuNP probe DNA | /5ThioMC6-D/AA AAA AAA AAA AAA ACC CGA GCA GCG CCC C |
| AuNP Capture DNA | /5AmMC6/AA AAA AAA AAA AAA AGG GGC GCT GCT CGG G |
| AR-10 | /5AmMC6/AA AAA AAA AAA AAA AAA AAA AAA AAC CCG ACC AGC CAC CAT CAG CAA CTC TTC CGC GTC CAT CCC TGC TG |
| HD4 | AAA AAA AAA AAA AAA AAA AAA GAT TTG TTT TCC CGA TTT TGC CCA GGG TTA ATT |

Table S2. DNA sequences used in this work.

**Surface chemistry process**

Gold nanoparticles-DNA conjugation: Lyophilized thiol-modified oligonucleotides (IDT) were suspended at a concentration of 500 µM in milliQ water; 50 µL of this solution was diluted 10x in 0.15 sodium phosphate buffer (pH 8.5) supplemented with 0.1 M DDT for 2 hours at room temperature to reduce the thiol group. A NAP 5 column with milliQ water as the mobile phase was used to elute the reduced oligonucleotide; concentration was approximately 25 µM. 0.1 mL of cytodiagnostics nanoparticles (of various sizes) were mixed with 20 µL of the purified DNA and 20 µL of 400 µg/mL thiol-modified PEG and allowed to incubate on a shaker at room temperature for 48 hours. The mixture was centrifuged at 0.4 rcf for 30 minutes and the supernatant was removed; the pellet was resuspended in 200 µL of TE buffer with 0.025% TWEEN-20 and the centrifugation and washing was repeated three times. Conjugated nanoparticles were stored in 4ºC.

GPTMS DNA surface modification: After GPTMS surface modification (see main text), 50 µM amine-modified DNA in milliQ water was incubated on the PC overnight at 4ºC. The surface was washed three times with syringe-filtered PBS (pore size 0.22 µm) before nanoparticles were added. Unbound nanoparticles were removed with filtered PBS.

Exosomes: PCs were first treated with 5% (3-Aminopropyl) tiethoxysilane (APTES) in tetrahydrofuran for 1 hour. Then a 10% dimethyl sulfide N,N'-Discuccinimidyl carbonate linker solution incubation was performed to functionalize the APTES for antibody binding for 45 minutes. The treated PCs were incubated with CD 147 antibody solution (0.1ug/uL) for 2 hours before exposing them to exosomes (7.5E7 particles/uL) retrieved from mouse plasma 2 hours. The sample is rinsed with PBS buffer to remove loose exosomes after incubation.

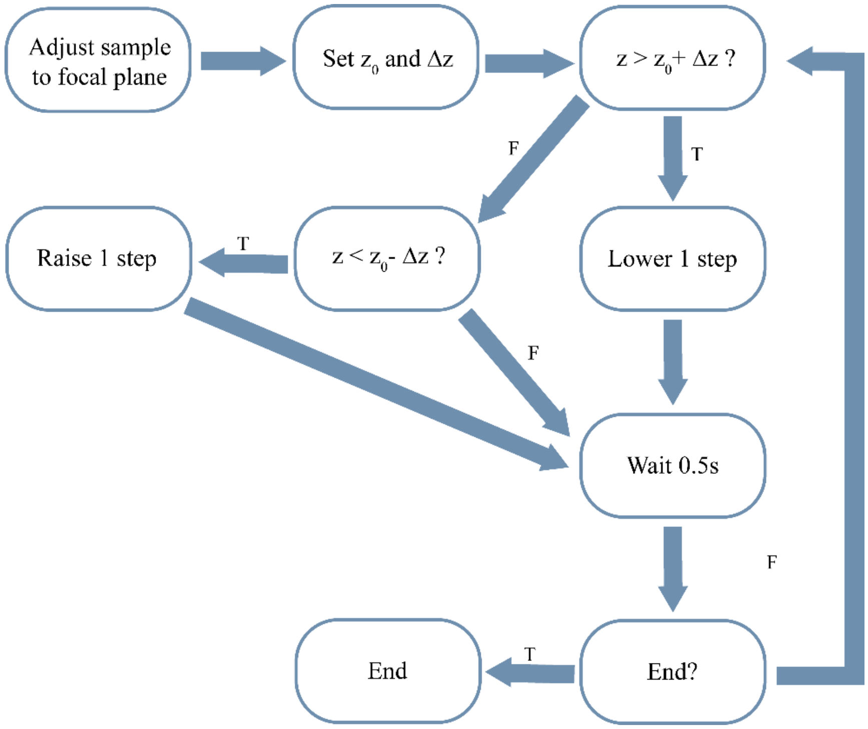


Fig. S1. Block diagram for AF algorithm.

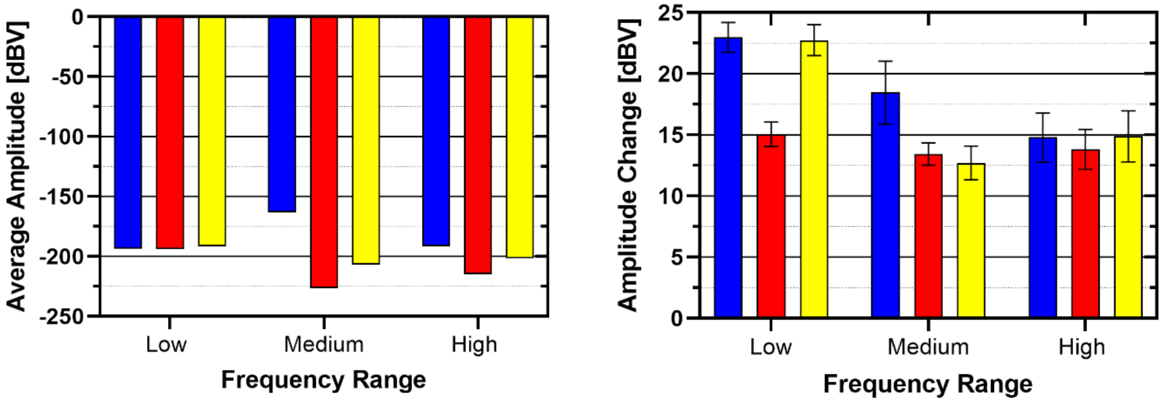


Fig. S2. (a) Average vibration amplitude comparison over three frequency ranges. Low: 1-10 Hz, Medium: 10-100 Hz, High: 100-1000 Hz. (b) Average change of vibration amplitude over ten human interference tests, where intentional stepping occurs near the instrument.

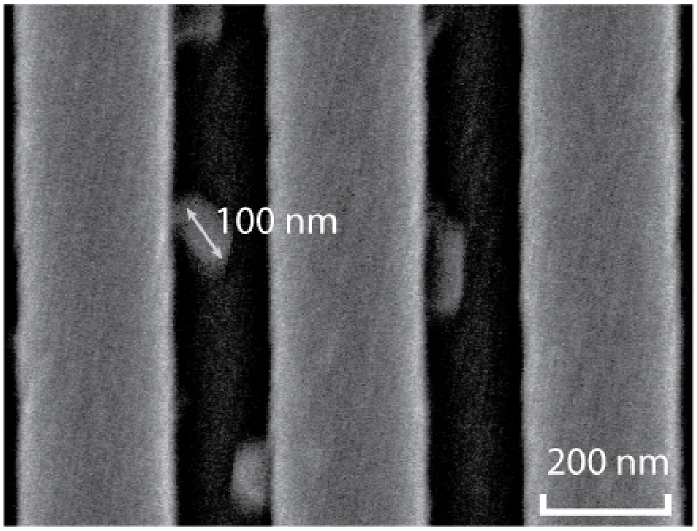


Fig. S3. SEM image of exosomes.

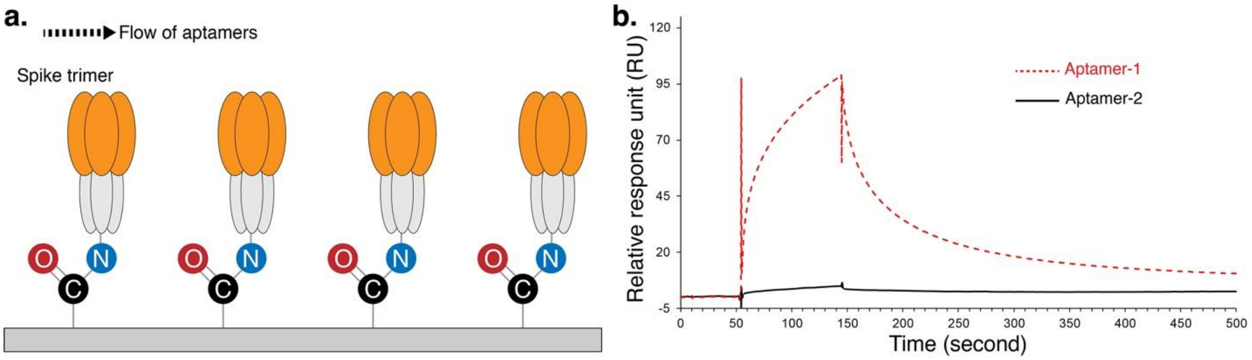


Fig. S4. (a) Schematic of the SPR assay used to determine the relative binding strength of RBD-1C aptamer (selected against spike RBD) and AR10 aptamer (selected against the entire pseudo-typed viral particle) to SARS-CoV-2 trimeric spike proteins. (b) Relative spike trimer binding response curves of RBD-1C aptamer (red dotted line) and AR10 aptamer (black line) show that AR10 does not interact with SARS-CoV-2 spike protein.

#### Surface plasmon resonance binding analysis procedures

Purified wildtype SARS-CoV-2 trimeric spike protein (Meridian Bioscience, Ohio, USA) was immobilized onto a research grade CM5 S-Series SPR chip (GE healthcare, Uppsala, Sweden) according to a standard amine coupling protocol. Briefly, carboxymethyl groups on the CM5 chip surface in Flow Cells 1 (as reference) or 2 were activated using a 420-second injection pulse at a flow rate 5 µL/min using a 4:1 mixture of N-ethyl-N-(dimethyaminopropyl) carbodiimide (EDC) and N-hydroxysuccinimide (NHS), respectively (final concentration of 200 mM EDC and 50 mM NHS, mixed immediately before injection). Following the activation, a 50 µg/mL Purified Mouse IgG (ImmunoReagents, North Carolina, USA) solution was prepared in a 10 mM sodium acetate (pH 5.0) buffer and then injected over the activated biosensor surface of Flow Cell 1. Following the activation, a 50 µg/mL SARS-CoV-2 trimeric spike protein was prepared in a 10 mM sodium acetate (pH 5.0) buffer and injected over the activated biosensor surface of Flow Cell 2. Excess unreacted carboxymethyl groups on the sensor surface were deactivated with a 600-second injection of 1 M ethanolamine in Flow Cells 1 or 2 at a flow rate 5 µL/min. Flow Cell 1 with immobilized Purified Mouse IgG served as a reference for Flow Cell 2 with immobilized trimeric spike protein. Different DNA aptamers were injected over the sensor chip at a flow rate of 5 µL/min with HBST-Mg, pH 7.4 (20 mM HEPES, 150 mM NaCl, 5 mM $MgCl_2$, 0.05% Tween-20) as running buffer. SPR measurements for relative response (RU) of different aptamers were performed on a BIAcore T200 (GE healthcare, Uppsala, Sweden) operated using the BIAcore T200 control software.

##### Explanation for the higher contrast signal of p-SARS-CoV-2

AR10 aptamer was selected against the entire SARS-CoV-2 virion. The SPR assays above show that AR10 does not bind to SARS-CoV-2 spike protein, suggesting that AR10 interacts with some moiety right on the viral membrane surface. Thus, the surface immobilized AR10 aptamers shall capture and bring the SARS-CoV-2 virions very close to the PC surface. On the contrary, HD4 aptamer was selected specifically to target the crown of gp120 protein, which is flexible and protruding from the HIV-1 viral membrane surface with a ~10 nm spacing. As a result, HIV-1 virions captured by the surface immobilized HD4 aptamers are farther from the PC surface than AR10 captured SARS-CoV-2 virions. The closer to the PC surface, the more enhancement is experienced by the pseudoviruses, leading to a higher contrast signal for the SARS-CoV-2 virions than the HIV virions.
